# Cell-type-specific astrocytic feedback regulates excitation–inhibition balance and cortical network dynamics

**DOI:** 10.64898/2026.09.26.754623

**Authors:** Den Whilrex Garcia, Sabir Jacquir

## Abstract

Astrocytes actively regulate synaptic transmission and neuronal excitability, yet their role in orchestrating macroscopic cortical network regimes and slow-wave oscillations remains an active area of reasearch. This study investigates how bidirectional neuron–astrocyte interactions shape emergent population dynamics using a computational network model of excitatory and inhibitory neurons coupled to an astrocyte. The results identify astrocytic feedback topology, rather than astrocytic coupling strength alone, as a key determinant of emergent cortical network dynamics. By systematically dissecting pathway-specific connectivity, it has been shown that the neuronal population driving astrocytic activation and the neuronal population receiving gliotransmission jointly determine whether the network occupies asynchronous irregular (AI), synchronous irregular (SI), synchronous regular (SR), asynchronous regular (AR) or quiescent regimes. Directing gliotransmission selectively onto excitatory neurons consistently promotes population synchrony regardless of the population influencing astrocytic dynamics, whereas selective modulation of inhibitory interneurons induces network quiescence via strong suppression. Under dual-target gliotransmission, network synchrony is dictated by the population driving astrocytic dynamics: excitatory-only drive promotes synchrony, while combined or inhibitory-specific drive preserves asynchronous states. Furthermore, the model reveals that astrocytic signaling kinetics provide an additional temporal control mechanism that regulates the frequency and persistence of self-sustained up states.

**Highlights:**

- Cell-type-specific astrocytic feedback governs emergent cortical dynamics and *E*–*I* balance.
- Astrocyte driven by excitatory neurons enforces synchrony, whereas inhibitory induces quiescence.
- Astrocyte signaling kinetics regulate self-sustained up states.

## 1. Introduction

Astrocytes are star-shaped glial cells that represent the most abundant non-neuronal cell type in the central nervous system (CNS) (Verkhratsky and Nedergaard, 2018; Chen et al., 2023; Stogsdill et al., 2023). Once regarded as passive support elements, astrocytes are now recognized as heterogeneous and dynamic regulators equipped with a rich repertoire of receptors, ion channels, transporters, and secretory pathways that enable them to sense and respond to local neural activity and metabolic demands (Verkhratsky and Nedergaard, 2018; Rupareliya et al., 2023). Their intricate, highly ramified processes tile the brain parenchyma with minimal spatial overlap (Bushong et al., 2002; Ogata and Kosaka, 2002), contacting synapses (Allen and Eroglu, 2017), cerebral vasculature (Zhou et al., 2019), and neighboring glia (Orthmann-Murphy et al., 2008) to form gap-junction-coupled syncytia spanning extensive tissue volumes (Cooper et al., 2026). Through this organization, astrocytes are integrated into neural circuits and participate in CNS homeostasis across molecular, network, and systems levels (Han et al., 2021; Lee et al., 2022).

Functionally, astrocytes execute diverse tasks essential for normal brain function. They regulate extracellular ion and neurotransmitter homeostasis by buffering potassium ions (K^+^) and clearing glutamate via high-affinity transporters, thereby preventing excitotoxicity and stabilizing baseline excitability (Lee et al., 2022; Chen et al., 2023). Astrocytes also provide metabolic support by mobilizing glycogen stores, supplying lactate and other substrates to active neurons, and coupling neuronal energy demands to local blood flow as core elements of the neuro-glio-vascular unit (Chen et al., 2023; Stogsdill et al., 2023). At tripartite synapses, comprising an astrocyte, presynaptic neuron, and postsynaptic neuron, they actively guide development and plasticity by modulating synaptogenesis, pruning connections, and releasing gliotransmitters (e.g., glutamate, ATP, and D-serine) alongside neurotrophic factors (Verkhratsky et al., 2016; Augusto-Oliveira et al., 2020). Furthermore, astrocytes serve as homeostatic sentinels by maintaining blood–brain barrier integrity and participating in neuroimmune signaling during physiological and reactive states (Lee et al., 2022; Fisher and Liddelow, 2024). Because astrocytes modulate synaptic transmission and circuit excitability on time scales slower than fast action potentials but faster than long-term structural remodeling, they are increasingly recognized as master modulators of neural networks (Augusto-Oliveira et al., 2020). Disruption of astrocytic signaling is implicated in diverse neurological and psychiatric disorders, underscoring the necessity of intact neuron–astrocyte interactions for cognitive function and network stability (Lee et al., 2022; Stogsdill et al., 2023; Valles et al., 2023). These insights motivate modeling frameworks that treat neuronal computation and astrocytic regulation as coupled, bidirectional processes rather than isolated systems.

This shift in perspective has driven increasing interest in computational models that explicitly incorporate astrocytes alongside neurons. At the cellular scale, computational models have demonstrated how intracellular calcium (Ca^2+^) dynamics and gliotransmitter release can regulate synaptic transmission and plasticity (Oschmann et al., 2018). Extending these formulations to the population level, neuron–astrocyte network models have been used to investigate synchronization, information processing, and the regulation of cortical network states (Manninen et al., 2023; Ali et al., 2025). Importantly, astrocytic modulation has also been implicated in the emergence and switching of cortical up and down states, suggesting that neuron–astrocyte interactions may contribute to the generation and stabilization of distinct network regimes (Blum Moyse and Berry, 2022). However, an important question remains unresolved: which features of neuron–astrocyte communication determine the direction and magnitude of these network-level effects? In particular, it remains unclear whether astrocytes primarily regulate global network dynamics through the strength of their coupling to neuronal populations or through the specific organization of the feedback pathways, including which neuronal population drives astrocytic activation and which population receives gliotransmission.

In this work, it is hypothesized that astrocytic feedback does not act as a uniform modulatory mechanism, but instead produces cell-type- and pathway-specific effects on cortical network dynamics. To test this hypothesis, a large recurrent spiking network comprising excitatory regular-spiking (RS) and inhibitory fast-spiking (FS) neurons coupled bidirectionally to a representative astrocytic signaling model was developed. This framework builds on previous computational studies of neuron–astrocyte interactions and incorporates adaptive exponential integrate-and-fire neuronal dynamics together with astrocytic Ca^2+^ signaling and gliotransmitter-mediated feedback (Brette and Gerstner, 2005; Postnov et al., 2007; Naud et al., 2008; Postnov et al., 2009; Garcia and Jacquir, 2024). A systematic variation of the neuronal populations driving astrocytic activation, the neuronal populations receiving astrocytic feedback, and the strength of cell-type-specific astrocytic coupling were performed. Through such approach, the effects of feedback topology, coupling strength, and connection probability on emergent network dynamics and whether specific neuron–astrocyte pathways preferentially promote synchrony, asynchronous activity, or network suppression were determined.

Beyond externally driven network states, it was further investigated whether astrocytic signaling can regulate the temporal organization of self-sustained activity following the removal of external input. Previous work has shown that astrocytic dynamics can contribute to the emergence of persistent neuronal activity and slow network oscillations (Garcia and Jacquir, 2024, 2025). Examination of how the astrocytic signaling parameters and cell-type-specific coupling influence both the frequency and persistence of network up states were therefore conducted. Together, these analyses provide a systematic characterization of how the cellular specificity, pathway topology, and temporal kinetics of astrocytic feedback shape emergent cortical network dynamics. Rather than considering astrocytes as homogeneous modulators of neuronal activity, the developed model predicts that their functional impact depends critically on which neuronal populations drive astrocytic activation and which populations receive astrocytic feedback. These results identify specific neuron–astrocyte pathways and astrocytic parameters that generate experimentally testable predictions regarding the regulation of cortical network states.

## 2. Methods

### 2.1. Neuron-astrocyte network model

Neuronal dynamics were described using the Adaptive Exponential Integrate-and-Fire (AdEx) model, defined by two coupled ordinary differential equations governing the membrane potential *V* and the adaptation current *w* (Brette and Gerstner, 2005; Touboul and Brette, 2008):

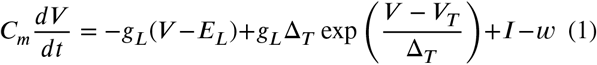

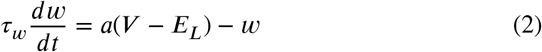

where *I* denotes the total input current, comprising excitatory and inhibitory conductance-based synaptic currents (*I* syn_*e*_ and *I* syn_*i*_) and an astrocytic current *I* ast that mimics the modulatory effects of gliotransmission. Here, *I* ast is modeled as glutamate-mediated current, as glutamate signaling represents the most well-characterized mechanism of astrocytic communication (Santello and Volterra, 2009; Garcia and Jacquir, 2024). The total input current is expressed as:

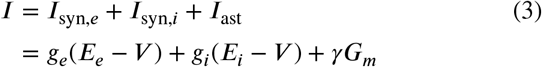

where *γ* ∈ [0, 1] represents the coupling strength between the astrocyte and the neuron (*γ* = 0 denotes no astrocytic influence), and *G*_*m*_ denotes the level of glial mediator production. The exponential term in Eq. 1 drives a rapid divergence in *V* upon sufficient depolarization, initiating a spike when a predefined threshold is crossed for computational purposes (Brette and Gerstner, 2005; Barranca et al., 2014). Following a spike, the membrane potential is reset to *V*_*r*_ and the adaptation current is incremented by *b* (Brette and Gerstner, 2005; Zerlaut et al., 2018):

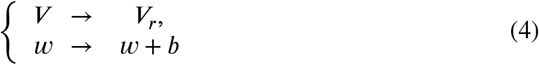

Chemical synapses are modeled as conductance-based connections, where each presynaptic spike induces an instantaneous step increase in the corresponding excitatory (*g*_*e*_) or inhibitory (*g*_*i*_) postsynaptic conductance, which subsequently decays exponentially over time (Stimberg et al., 2019b):

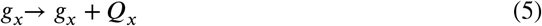

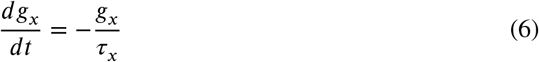

where *x* ∈ [*e, i*] denotes excitatory or inhibitory synapses, *Q*_*x*_ is the conductance increment per presynaptic event, and τ_*x*_ is the corresponding synaptic time constant.

Meanwhile, astrocytic Ca^2+^ dynamics are described using a set of ordinary differential equations adapted from the reduced framework of prior studies (Postnov et al., 2007, 2009):

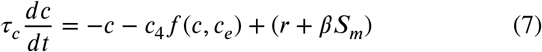

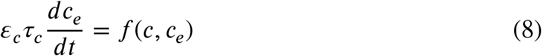

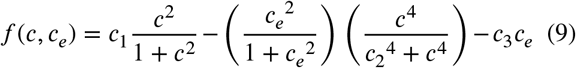

Here, *c* and *c*_*e*_ denote the cytosolic and endoplasmic reticulum (ER) Ca^2+^ concentrations, respectively, with the ER serving as the primary intracellular store. The nonlinear exchange between these two compartments is captured by the function *f* (*c, c*_*e*_), while τ_*c*_ sets the characteristic timescale of Ca^2+^ oscillations and transients. Additionally, *S*_*m*_ represents a secondary mediator production regulated by the factor *β*. Although this formulation adopts a deterministic description—thus omitting local and global stochastic effects—it retains sufficient complexity to reproduce the key dynamical features of astrocytic Ca^2+^ signaling (Garcia and Jacquir, 2024).

The variables *S*_*m*_ and *G*_*m*_ represent the secondary messenger and gliotransmitter mediator production, respectively. Following the simplified event-driven scheme proposed in recent work (Garcia and Jacquir, 2024)—based on the original formulation of previous studies (Postnov et al., 2007, 2009)—their dynamics are driven by discrete step increases triggered by specific events, followed by exponential decay. Specifically, *S*_*m*_ is updated upon each presynaptic spike, whereas *G*_*m*_ is triggered when cytosolic Ca^2+^ crosses a critical threshold. In general, these kinetics can be expressed as:

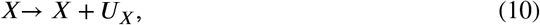

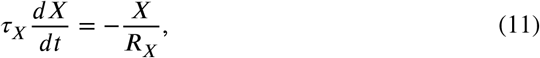

where *X* ∈ [*S*_*m*_, *G*_*m*_]. Here, *U*_*X*_ denotes the event-dependent increment, τ_*X*_ the characteristic timescale, and *R*_*X*_ the decay parameter. Complete parameter definitions and values are listed in Table 1.

**Table 1.** Model parameter values for the neurons, astrocytes, synapses, and messenger kinetics (Postnov et al., 2007; Naud et al., 2008; Postnov et al., 2009; Garcia and Jacquir, 2024, 2025).

| Parameter | Description | Values |  |
| --- | --- | --- | --- |
| *Neuron Model Parameters* |  | *Exc* | *Inh* |
| $C_m$ | Membrane capacitance | 200 pF | 200 pF |
| $g_L$ | Leak conductance | 10 nS | 10 nS |
| $V_T$ | Effective threshold potential | −50 mV | −50 mV |
| $V_r$ | Reset potential | −65 mV | −65 mV |
| $\tau_w$ | Adaptation time constant | 500 s | 500 s |
| $a$ | Subthreshold adaptation | 0 nS | 0 nS |
| $b$ | Spike-triggered adaptation increment | 10 pA | 0 pA |
| $\Delta_T$ | Threshold slope factor | 2 mV | 0.5 mV |
| $E_L$ | Leak reversal potential | −66 mV | −67 mV |
| $V_{cut}$ | Firing threshold | −40 mV | −47.5 mV |
| *Synapse Model Parameters* |  | *Exc* | *Inh* |
| $E_x$ | Synaptic reversal potential | 0 mV | −80 mV |
| $\tau_x$ | Synaptic time constant | 5.5 ms | 5.0 ms |
| $Q_x$ | Synaptic conductance increment | 1.5 nS | 5.0 nS |
| *Astrocyte Model Parameters* |  |  |  |
| $\epsilon_c$ | Time separation parameter | | 0.04 |
| $r$ | Initial state of cytosolic $\text{Ca}^{2+}$ concentration | | 0.31 |
| $c_{peak}$ | Cytosolic $\text{Ca}^{2+}$ concentration threshold | | 0.6 |
| $c_1$ | Control parameter 1 for cytosolic $\text{Ca}^{2+}$ concentration | | 0.13 |
| $c_2$ | Control parameter 2 for cytosolic $\text{Ca}^{2+}$ concentration | | 0.9 |
| $c_3$ | Control parameter 3 for cytosolic $\text{Ca}^{2+}$ concentration | | 0.004 |
| $c_4$ | Control parameter 4 for cytosolic $\text{Ca}^{2+}$ concentration | | $2/\epsilon_c$ |
| *Messenger Production Model Parameters* |  |  |  |
| $R_{S_m}$ | $S_m$ decay parameter | | 3 |
| $R_{G_m}$ | $G_m$ decay parameter | | 3 |
| $U_{S_m}$ | $S_m$ increment parameter | | 0.6 |
| $U_{G_m}$ | $G_m$ increment parameter | | 0.6 |

### 2.2. Spiking network configuration

In the present work, the network comprised 10,000 neurons, including 8,000 regular-spiking (RS) excitatory neurons and 2,000 fast-spiking (FS) inhibitory neurons, consistent with the commonly used 80:20 excitatory:inhibitory ratio reported for cortical networks and with the established association of RS and FS firing phenotypes with excitatory and inhibitory cell classes, respectively (Kilgore et al., 2025; Soriano et al., 2008; Connors and Gutnick, 1990). Furthermore, following previous modeling works (Lorenzo et al., 2020; Vuillaume et al., 2021; Tesler et al., 2023; Lorenzo et al., 2025), local neuron–glial interactions are abstracted here by coupling a single, representative astrocyte to the surrounding neuronal population. This modeling choice aligns with physiological findings showing that *in vivo* Ca^2+^ dynamics tend to be compartmentalized within individual astrocytic domains (Halassa et al., 2007; Schummers et al., 2008; Breithausen et al., 2020). Consequently, while the present model bypasses the extended syncytial networks and propagating intercellular calcium waves typical of *in vitro* preparations (Guthrie et al., 1999), it provides an efficient and biologically grounded description of single-cell astrocytic modulation.

Each neuronal population receives driving input from an external Poisson group equal in size to the excitatory population. The connection probability from this Poisson source to both excitatory and inhibitory populations is fixed at 0.05, matching the recurrent connectivity within the neuronal network. The external drive was applied for 5s, while the total simulation duration was 20s, enabling the assessment of whether the network can maintain self-sustained activity after the removal of external stimulation. When astrocytic modulation is included, baseline parameters were set as follows unless stated otherwise: characteristic timescale of astrocytic Ca^2+^ dynamics τ_*c*_ = 8ms, decay time constant of the secondary messenger 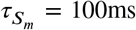, decay time constant of the gliotransmitter mediator 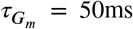, control parameter of the slow activation pathway *β* = 0.50, and coupling strength to excitatory neurons *γ*_*e*_ = 0.50 (Postnov et al., 2007, 2009). Directional connectivity between the astrocyte and the neuronal population was established with a neuron-to-astrocyte probability of *prb*_*NA*_ = 0.25 and an astrocyte-to-neuron feedback probability of *prb*_*AN*_ = 0.05 (Blum Moyse and Berry, 2022). To isolate their specific effects, single-parameter sweeps were conducted for *β, prb*_*NA*_, *prb*_*AN*_, and the cell-type-specific coupling weights (*γ*_*e*_ and *γ*_*i*_, bound by *γ*_*e*_ + *γ*_*i*_ = 1), while keeping all remaining parameters fixed at baseline values.

A schematic of the network configuration and neuron–astrocyte interactions is shown in Figure 1. All simulations of the neuron–astrocyte network were executed using the Brian2 spiking neural network simulator in Python (Stimberg et al., 2019a).

**Figure 1.**
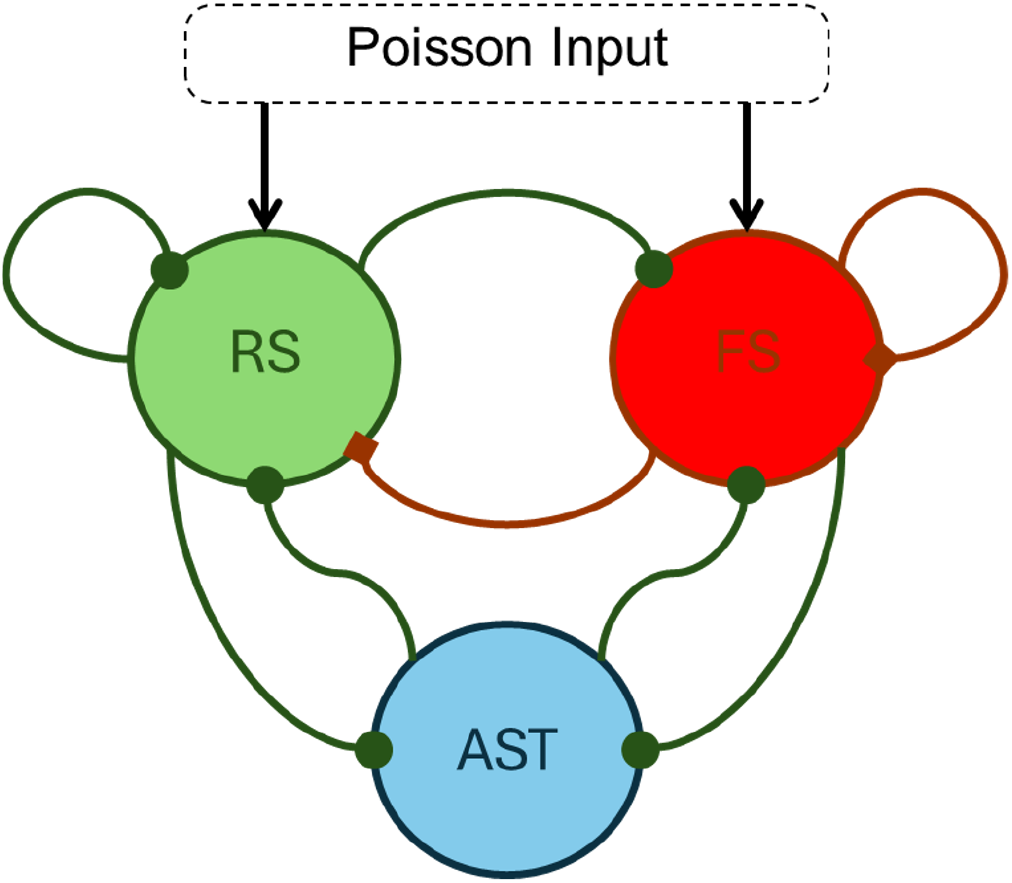
Schematic of the network configuration. Green circles represent RS excitatory neurons, red circles denote FS inhibitory neurons, and blue represents the astrocyte. Green arrows terminating in circles indicate excitatory synapses, while red arrows ending in diamonds indicate inhibitory synapses. Both neuronal populations receive input from a Poisson source. Seed number 7 was used to ensure reproducibility.

### 2.3. Evaluation of Network Dynamics and State Transitions

Quantification of network dynamics relied on two distinct metrics: spiking regularity, which evaluates temporal activity patterns, and population synchrony, which measures firing coordination across the neuronal ensemble.

Spiking regularity was quantified using the coefficient of variation (CV) of interspike intervals (ISI), calculated across a representative subset of the network:

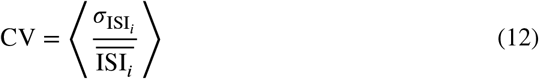

where 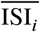 and 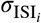 denote the mean and standard deviation of the interspike intervals for neuron *i*, respectively. The operator ⟨⋅⟩ represents the population average over 10% of the network, selected via uniform random sampling without replacement. Values of CV ≥ 1 reflect temporally irregular spiking and were used here to classify irregular network activity.

Population synchrony was quantified through the average pairwise spike-count cross-correlation (CC) across the network:

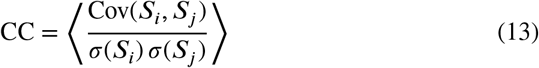

where Cov(*S*_*i*_, *S*_*j*_) represents the covariance between the binned spike counts *S*_*i*_ and *S*_*j*_ of neurons *i* and *j, σ*(⋅) denotes the standard deviation, and ⟨⋅⟩ indicates the ensemble average over sampled pairs. By default, spike trains were discretized into 5ms bins, and correlations were computed across 1000 randomly selected, disjoint pairs (800 excitatory and 200 inhibitory). The cross-correlation index ranges from −1 to 1, with higher positive values indicating synchronous firing, while lower values demonstrating asynchronous firing. These conditions for spiking regularity and population synchrony are consistent with prior criteria (Destexhe, 2009). It is also important to note that if a neuron exhibited insufficient spiking activity or zero spike-count variance across bins, the value can be undefined; these pairs were omitted from the population average.

In addition, up and down states were identified from the excitatory population firing rate using a threshold-based detection algorithm. Spike trains were binned at a temporal resolution of 1ms and smoothed with a Gaussian filter to suppress high-frequency fluctuations. Periods during which the smoothed population rate exceeded 20% of its maximum value were provisionally designated as up states. To eliminate spurious detections, only sustained up states lasting longer than 100ms were retained; shorter ones were either merged with the preceding state or discarded. This detection pipeline was applied across the entire simulation duration to evaluate self-sustained dynamics, where the persistence of up states after the removal of external input signified self-sustained network activity. Figure 2 illustrates a representative example of how up and down states were detected.

**Figure 2.**
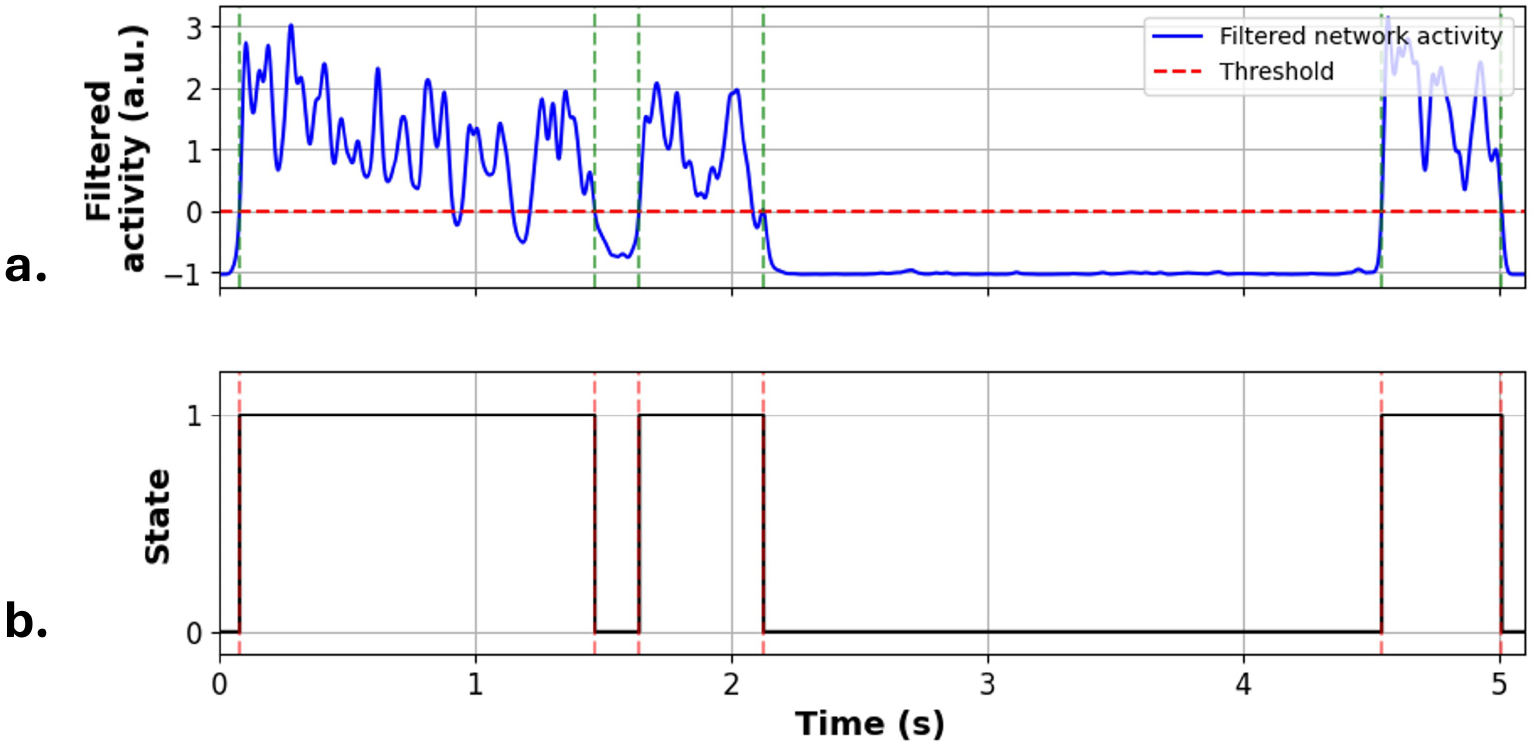
Detection of up and down states from population ring dynamics. (a) Filtered population ring rate (blue trace) alongside the detection threshold set at 20% of peak network activity (horizontal red line). Periods sustained above threshold for at least 100ms are identified as up states, with state transition boundaries marked by vertical green lines. (b) The black trace displays the resulting binary network states obtained from the smoothed excitatory ring rate (1ms bins with Gaussian filtering), where 1 corresponds to an Up state and 0 to a Down state.

## 3. Results and Discussion

### 3.1. Astrocytic feedback reorganizes cortical network states

The initial analysis examined whether astrocytic modulation alters the regularity, synchrony, or both, of network dynamics. To evaluate this, the coefficient of variation (CV) and cross-correlation (CC) were computed exclusively during the external drive interval (0–5 s). Based on these metrics, four distinct network regimes were identified: synchronous irregular (SI; CV ≥ 1.0, CC ≥ 0.01), asynchronous irregular (AI; CV ≥ 1.0, CC < 0.01), synchronous regular (SR; CV < 1.0, CC ≥ 0.01), and asynchronous regular (AR; CV < 1.0, CC < 0.01). Typically, up and down states correspond to the SR regime, whereas wakefulness is generally associated with the AI regime (Destexhe, 2009; Torao-Angosto et al., 2021; Sanchez-Vives et al., 2025). Figure 3 maps the observed dynamical states across the simulated parameter space. Among the 1881 simulated network configurations, 502 cases exhibited minimal spiking activity, causing the cross-correlation (CC) to be undefined. These instances occurred when the astrocyte modulated only the inhibitory population (*γ*_*i*_ = 1.0), where strong inhibition suppressed network activity.

**Figure 3.**
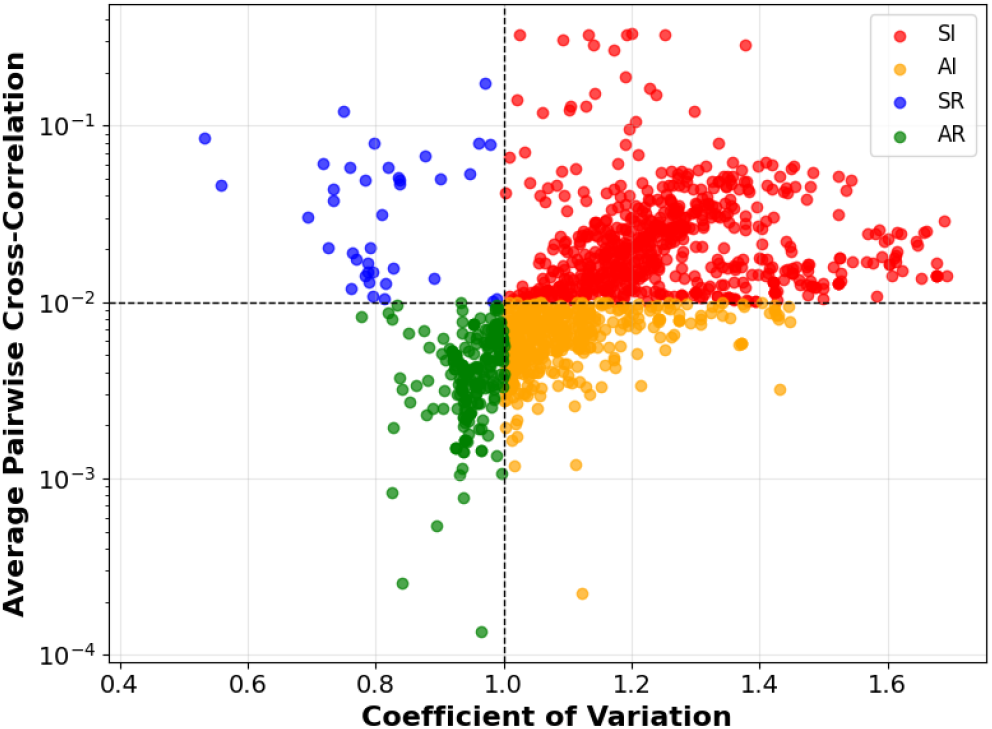
Classification of network activity regimes based on the coe-cient of variation (CV) of neuronal firing and pairwise cross-correlation (CC). Each dot represents a combination of CV and CC values for a given network configuration. Regimes include: SI (synchronous irregular) in red, AI (asynchronous irregular) in orange, SR (synchronous regular) in blue, and AR (asynchronous regular) in green.

Having established that four distinct regimes emerge, we next asked which feature of astrocytic feedback, the driving population, the target population, or both, determines which regime the network occupies.

### 3.2. Cell-type-specific feedback regulates excitation–inhibition balance

Among these, the relative astrocytic coupling weights onto excitatory and inhibitory populations (*γ*_*e*_ and *γ*_*i*_, constrained by *γ*_*e*_+*γ*_*i*_ = 1) emerged as critical drivers of network state transitions. Specifically, when astrocytic activation was driven exclusively by the inhibitory population while gliotransmitter release modulated both populations, the network displayed asynchronous dynamics at intermediate values of *γ*_*e*_, but shifted to synchronous regimes at both low and high *γ*_*e*_ limits (Figure 4a–c). Conversely, when both neuronal populations influenced the astrocyte but glial feedback targeted only excitatory neurons, the network remained asynchronous at low *γ*_*e*_ and transitioned monotonically into synchrony as *γ*_*e*_ increased (Figure 4d–f).

**Figure 4.**
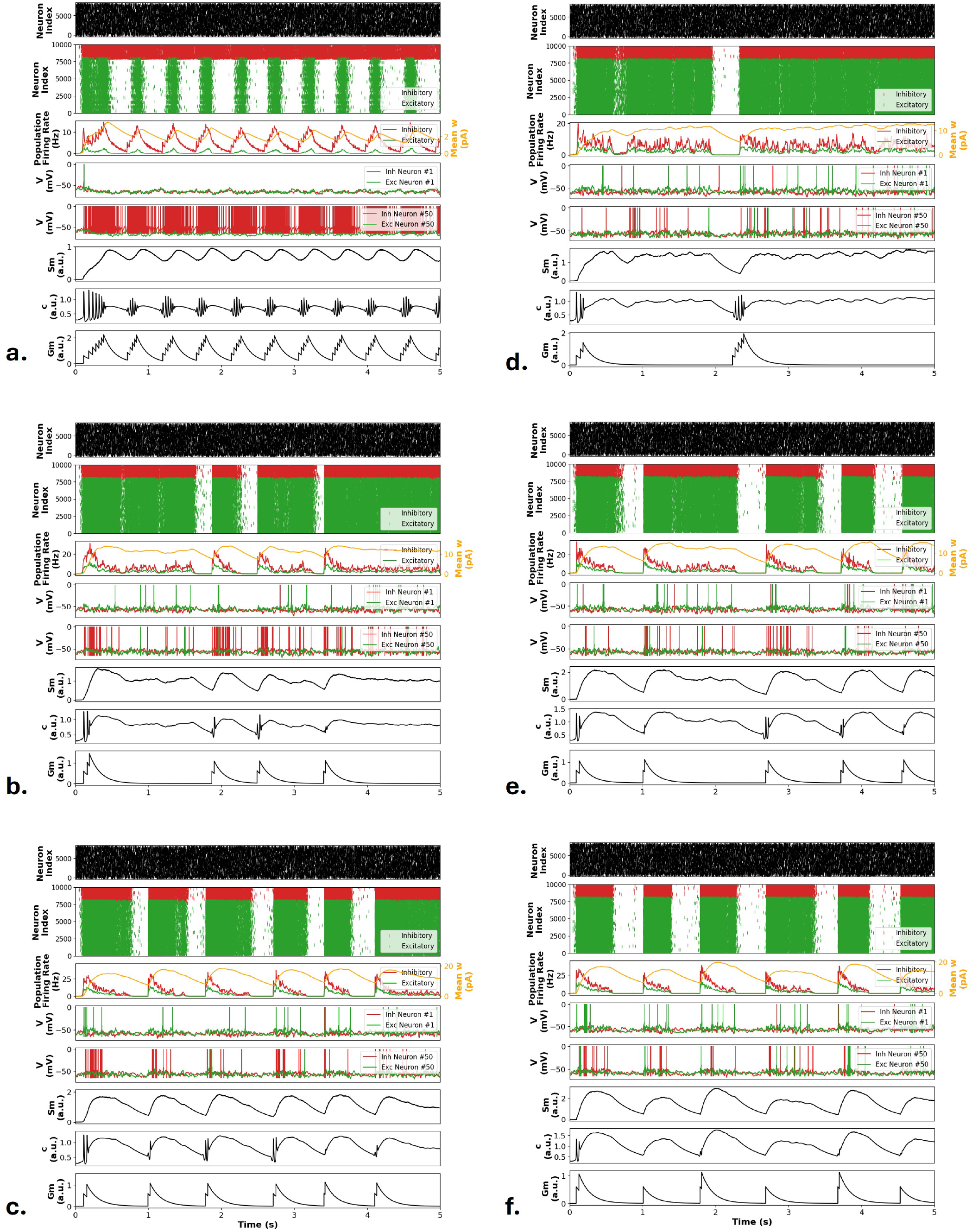
Network dynamics under varying neuron astrocyte coupling strengths (*γe*). (**a–c**) Astrocyte driven exclusively by inhibitory neurons with feedback to both populations (*γe* = 0.3, 0.6, 0.85). (**d–f**) Astrocyte driven by both populations with feedback targeted solely to excitatory neurons (*γe* = 0.05, 0.5, 0.95). Traces from top to bottom in each panel: Poisson input rasters; neuronal population rasters; mean E/I ring rates; adaptation currents and membrane potentials of representative neurons; and astrocytic secondary messenger (*Sm*), Ca^2+^, and gliotransmitter (*Gm*) dynamics.

The principal findings across these coupling-strength configurations are tabulated in Table 2. When astrocytic modulation was directed exclusively toward the inhibitory population, the network remained largely quiescent, as potentiated interneuron activity suppressed overall firing. Conversely, when glial feedback selectively targeted the excitatory population, the network shifted from asynchronous to synchronous dynamics as the excitatory coupling weight (*γ*_*e*_) increased, indicating that elevated excitatory drive disrupts the excitation–inhibition (E/I) balance to promote collective synchronization. When both populations received astrocytic input, asynchronous dynamics were preserved only within an intermediate range of *γ*_*e*_, where balanced E/I interactions were maintained; deviations toward either low or high coupling extremes disrupted this equilibrium and precipitated transitions into synchrony. Collectively, these findings highlight astrocyte–neuron coupling strength as a decisive parameter in regulating emergent population synchrony.

**Table 2.** Summary of network dynamics across neuron–astrocyte coupling strength configurations.

| Population Activating Astrocyte | Population Modulated by Astrocyte |  |  |
| --- | --- | --- | --- |
|  | Excitatory only | Inhibitory only | Excitatory + Inhibitory |
| **Excitatory only** | Predominantly asynchronous | Predominantly minimal to no activity | Synchronous at low $\gamma_e$; asynchronous at high $\gamma_e$ |
| **Inhibitory only** | Asynchronous at low $\gamma_e$; synchronous at high $\gamma_e$ | Predominantly minimal to no activity | Asynchronous at intermediate $\gamma_e$; synchronous at low and high |
| **Excitatory + Inhibitory** | Asynchronous at low $\gamma_e$; synchronous at high $\gamma_e$ | Predominantly minimal to no activity | Asynchronous at intermediate $\gamma_e$; synchronous at low and high |

Because coupling weight alone could not fully account for these transitions, it was next tested whether the topology of neuron–astrocyte connectivity independent of coupling strength was itself a determinant of network state.

### 3.3. Feedback topology determines the direction of astroctic modulation

Pathway topology acts as an all-or-none switch on network state, whereas coupling strength nitude of an already-active pathway — we tested this by systematically varying neuron-to-astrocyte and astrocyte-to-neuron connection probabilities under selective pathway activation.

Figure 5 was obtained by systematically varying the connection probabilities between the neuronal population and the representative astrocyte while selectively enabling or disabling specific signaling pathways. First, when astrocytic feedback was restricted exclusively to the excitatory population, synchronous regimes dominated regardless of whether astrocytic dynamics were driven by excitatory neurons, inhibitory neurons, or both, although asynchronous dynamics occasionally emerged at low neuron-to-astrocyte connection probabilities (leftmost columns of Figure 5). Conversely, when astrocytic modulation was directed only to the inhibitory population, the network consistently exhibited minimal activity or was completely silenced, as potentiated inhibition suppressed global spiking (middle column of Figure 5). Finally, when gliotransmission targeted both neuronal populations, network behavior depended heavily on the driving source: when either both populations or only the inhibitory population drove astrocytic dynamics, the network remained predominantly in an asynchronous state; in contrast, when only the excitatory population drove astrocytic dynamics, the network shifted primarily into synchrony (rightmost column of Figure 5). Together, these findings corroborate that the target of astrocytic modulation emerged as a major determinant of emergent network dynamics. This result suggests that the physiological consequence of astrocytic signaling may depend strongly on which neuronal population is embedded within the astrocyte-mediated feedback loop. While varying connection probabilities exerted comparatively modest effects, the active pathway topology played a decisive role. In contrast, systematic variations of other parameters—including *β*, τ_*c*_, τ_*Sm*_, and τ_*Gm*_—revealed no clear or systematic trends in network dynamics within the tested ranges, showing ambiguous effects on both synchrony and regularity.

**Figure 5.**
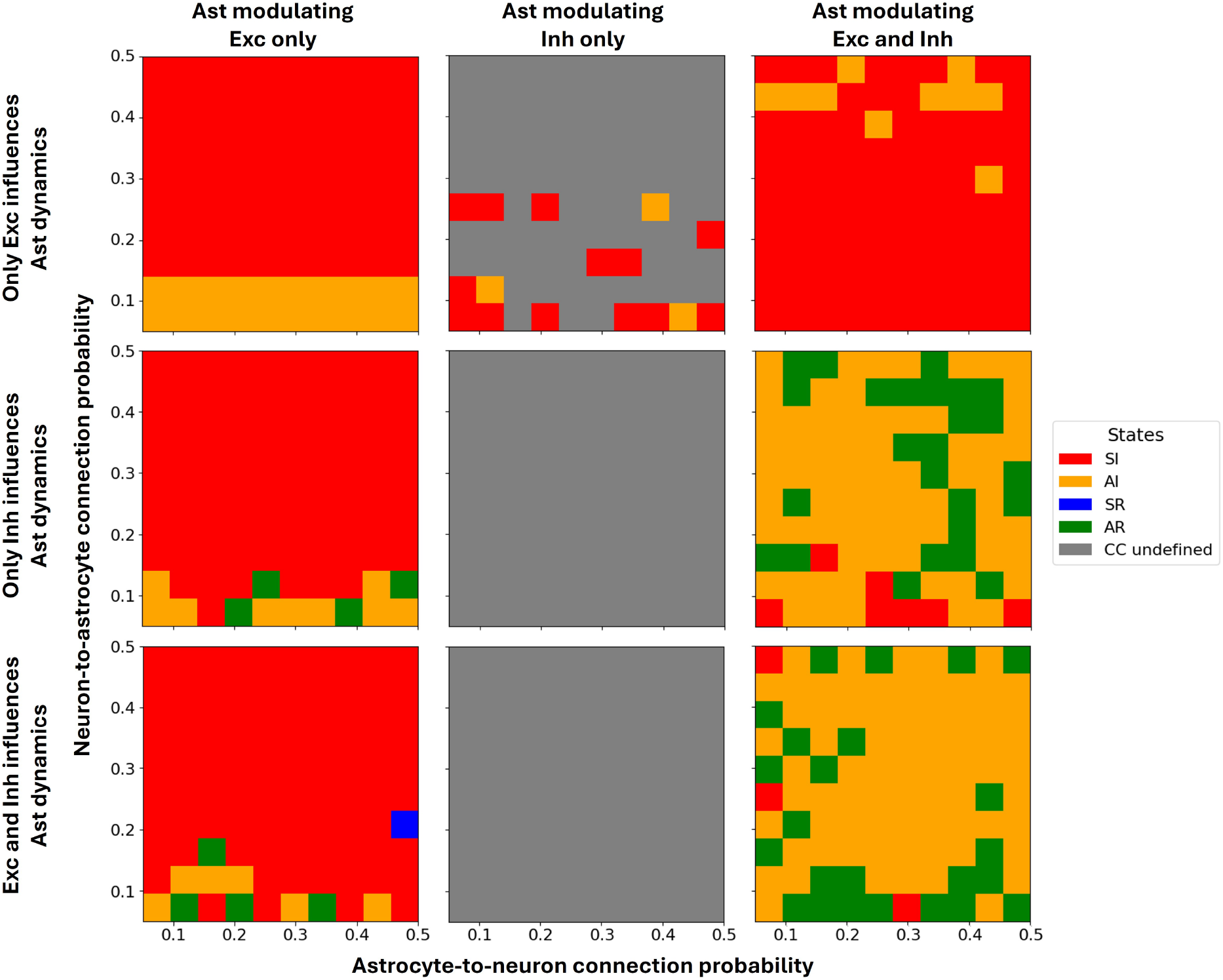
Network dynamical regimes across neuron–astrocyte connectivity configurations. Panels depict emergent network states as a function of neuron-to-astrocyte (vertical axis, *pr^b^_AN_*) and astrocyte-to-neuron (horizontal axis, *pr^b^_NA_*) connection probabilities under selective pathway activation. Rows indicate the driving neuronal population, while columns specify the target population modulated by gliotransmission. Color-coded regimes: synchronous irregular (SI, red), asynchronous irregular (AI, orange), synchronous regular (SR, blue), asynchronous regular (AR, green), and minimal to no activity/undefined cross-correlation (gray).

Beyond externally driven network states, it was examined whether the same astrocytic properties also shape the temporal organization of activity once external drive is removed.

### 3.4. Astrocytic signaling kinetics regulate the temporal persistence of up states

Having established that astrocytic feedback topology strongly influences the qualitative regime of network activity during external stimulation, it was next examined whether astrocytic signaling parameters also regulate the temporal organization of activity after stimulation is removed. This distinction is important because a parameter may influence the persistence or timing of collective activity without necessarily changing the qualitative dynamical regime of the network. Network configurations were evaluated to determine which could sustain oscillatory activity (self-sustained oscillations) following the removal of the external Poisson drive. Self-sustained dynamics were quantified by measuring the frequency of up states between 5–20s across various conditions (as shown in Figure 6). Up state counts exhibited an inverse relationship with the parameter *β*, systematically decreasing as *β* increased regardless of the driving neuronal population (Figure 6a). In contrast, up state generation remained largely robust to variations in the excitatory coupling strength *γ*_*e*_ (Figure 6b). Finally, increasing the time constants τ_*c*_, τ_*Sm*_, and τ_*Gm*_ consistently led to a reduction in up state frequency (Figure 6c–e). These results indicate that astrocytic signaling parameters regulate the temporal organization of recurrent network activity rather than simply determining whether the network is active or inactive. In particular, changes in *β* and in the characteristic time constants of astrocytic calcium and gliotransmitter dynamics altered the rate at which recurrent Up states emerged after stimulus removal. Thus, astrocytic dynamics may provide a temporal control layer that operates on top of the network-state transitions produced by feedback topology.

**Figure 6.**
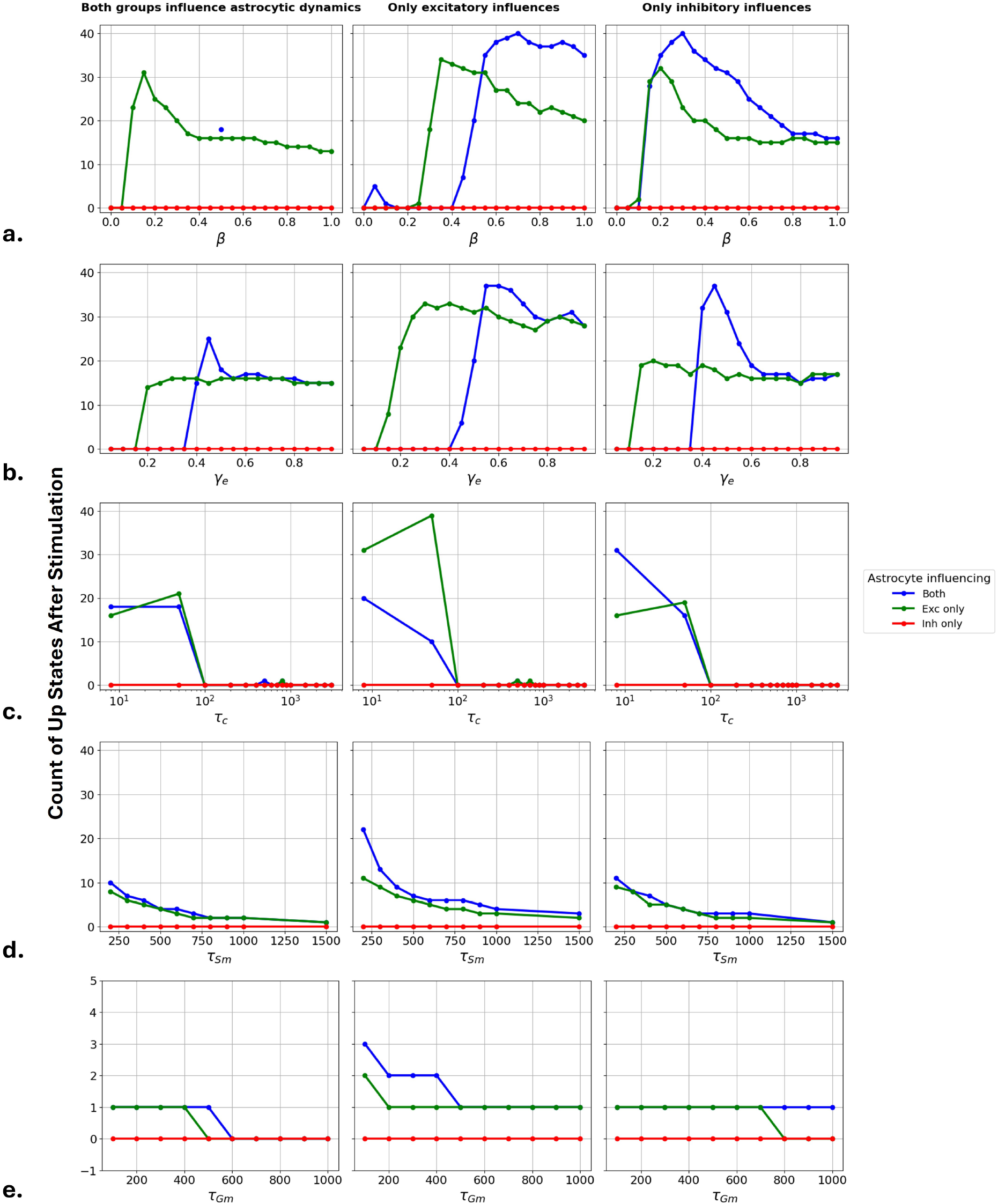
Total up state counts were recorded within the post-stimulation window. Columns (left to right) specify the neuronal population driving astrocytic dynamics: both (excitatory and inhibitory), excitatory only, and inhibitory only. Panels (**a–e**) illustrate parameter sweeps for *β, γ*_*e*_, τ_*c*_, τ_*Sm*_, and τ_*Gm*_, respectively. Line colors denote the target population receiving gliotransmission: both populations (blue), excitatory only (green), and inhibitory only (red).

To further evaluate the temporal characteristics of self-sustained dynamics, the mean duration of up states was quantified across parameter regimes (as shown in Figure 7). Low values of *β* sustained relatively long, stable up states that transitioned to shorter, plateaued durations as *β* increased (Figure 7a). A comparable dependency emerged for *γ*_*e*_, particularly when astrocytic dynamics were driven exclusively by excitatory neurons; in contrast, under both- or inhibitory-specific drive, mean up-state duration remained essentially invariant across *γ*_*e*_ levels (Figure 7b). Increasing the Ca^2+^ time constant τ_*c*_ systematically truncated up state duration, highlighting a pronounced sensitivity of network persistence to astrocytic calcium kinetics and suggesting that excessive values ultimately drive the network toward quiescence (Figure 7c). While variations in τ_*Sm*_ exerted negligible influence on state duration (Figure 7d), τ_*Gm*_ exhibited a biphasic effect, initially prolonging up states before causing a subsequent decline at higher values—except when gliotransmission targeted both populations, where duration scaled monotonically with τ_*Gm*_ (Figure 7e). Across all configurations, selective astrocytic modulation of the inhibitory population sustained profound suppression, preventing the emergence of post-stimulation up states.

**Figure 7.**
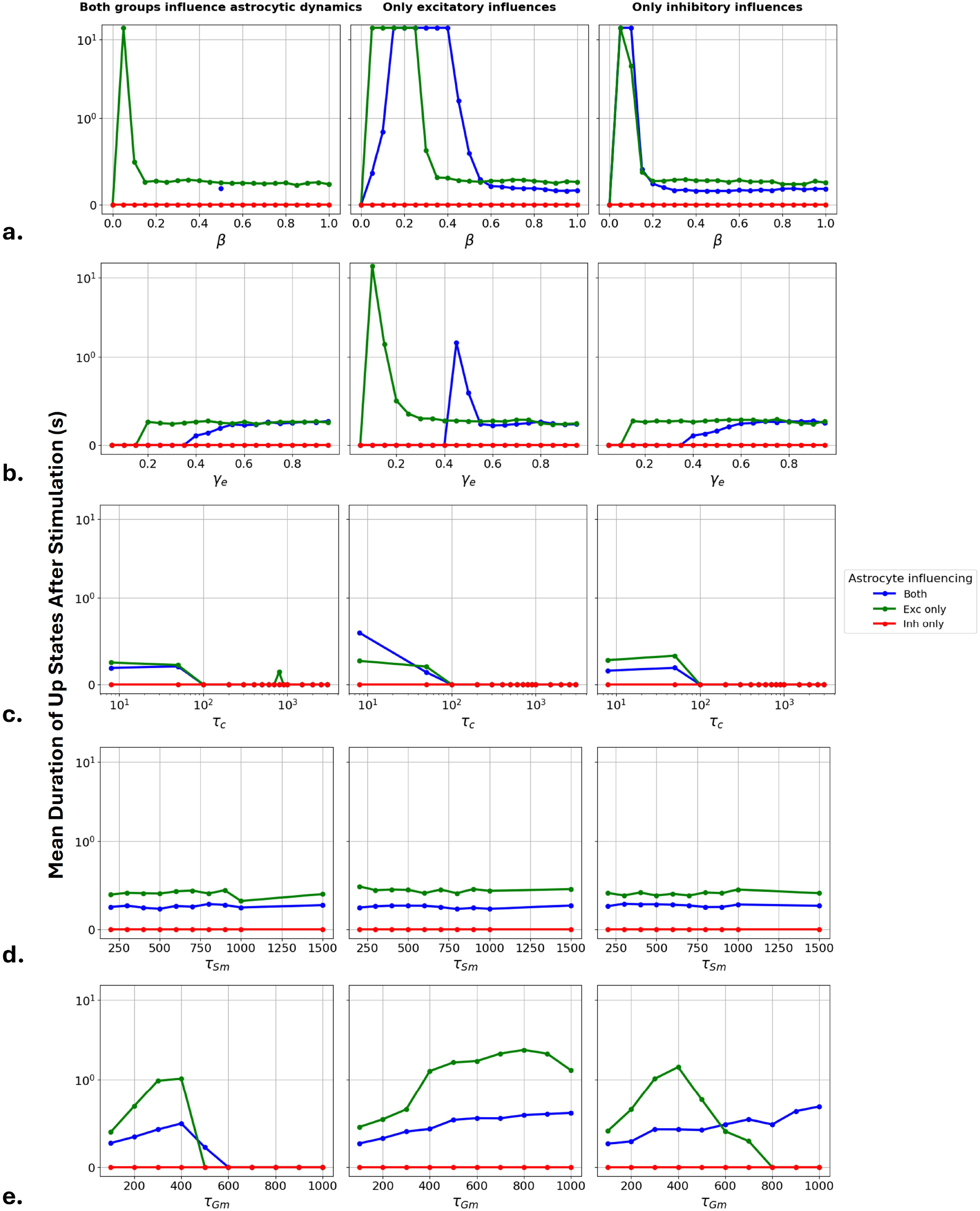
Mean duration of up states was recorded within the post-stimulation window. Columns (left to right) specify the neuronal population driving astrocytic dynamics: both (excitatory and inhibitory), excitatory only, and inhibitory only. Panels (**a–e**) illustrate parameter sweeps for *β, γe*, τ*c*, τ*Sm*, and τ*Gm*, respectively. Line colors denote the target population receiving gliotransmission: both populations (blue), excitatory only (green), and inhibitory only (red).

Notably, the same kinetic parameters (*β*, τ_*c*_, τ_*Sm*_, and τ_*Gm*_) that showed no systematic effect on network regime during external drive (Section 3.3) exerted strong, systematic control over the frequency and persistence of self-sustained activity — indicating that astrocytic kinetics govern the temporal, rather than the categorical, organization of network dynamics. Taken together, the frequency and duration analyses indicate that astrocytic signaling can regulate distinct temporal properties of self-sustained activity. Changes in astrocytic kinetics affected both the recurrence rate and persistence of up states, although the magnitude and direction of these effects depended on the signaling pathway and target neuronal population. This suggests that astrocytic feedback may provide multiple temporal control mechanisms for shaping collective cortical activity.

## 4. Conclusion

This study provides a systematic computational analysis of how cell-type-specific neuron–astrocyte interactions shape emergent cortical network dynamics. Rather than acting as a uniform modulatory mechanism, astrocytic feedback produced qualitatively different network effects depending on the neuronal populations driving astrocytic activation and receiving gliotransmission. Across the simulated parameter space, the target of astrocytic modulation emerged as a major determinant of network state: selective feedback to excitatory neurons promoted synchronous activity, whereas selective modulation of inhibitory neurons strongly suppressed network activity. When both neuronal populations were targeted, the network response depended on the relative strength of excitatory and inhibitory astrocytic coupling, revealing a parameter range in which balanced feedback supported asynchronous dynamics. These results identify the input–output organization of astrocytic signaling as an important determinant of emergent network synchrony and excitation–inhibition balance.

The model further showed that astrocytic feedback topology could have a stronger qualitative influence on network dynamics than variations in connection probability alone. In particular, changing which neuronal population activated the astrocyte and which population received astrocytic feedback could switch the network between asynchronous, synchronous, and quiescent regimes. This finding suggests that the functional impact of astrocytes may depend not only on the magnitude of gliotransmission, but also on the specific cellular pathways through which astrocytic signals are integrated into neuronal circuits. In this framework, astrocytic signaling provides a potential mechanism for cell-type-specific regulation of collective cortical activity.

From a physiological perspective, these findings suggest that astrocytic modulation may depend critically on the organization of local neuronal circuits rather than solely on the magnitude of gliotransmitter release. In particular, the neuronal population providing the dominant input to an astrocyte and the population receiving its feedback may determine whether astrocytic signaling reinforces recurrent excitation, enhances inhibitory control, or preserves excitation–inhibition balance. Selective astrocytic modulation of excitatory neurons may therefore provide a mechanism for amplifying coordinated population activity, whereas preferential modulation of inhibitory neurons may favor network suppression. Importantly, these interpretations should be viewed as testable mechanistic hypotheses rather than direct demonstrations of physiological circuit organization. In experimental settings, cell-type-specific manipulation of neuronal populations coupled to measurements of astrocytic activity could provide a means of testing whether distinct astrocyte–neuron pathways differentially regulate cortical synchrony and network state transitions.

The present model deliberately abstracts the spatial organization of astrocytic domains by representing a local astrocytic unit as a single dynamical element. This simplification allows the causal effects of cell-type-specific feedback pathways to be isolated, but it does not capture the spatial heterogeneity of astrocytic processes, intercellular calcium signaling, or astrocytic gap-junction coupling. Consequently, the predicted effects should be interpreted primarily as consequences of feedback organization rather than as quantitative representations of a specific cortical astrocytic microcircuit. Extending the framework to multiple spatially distributed astrocytes will be important for determining how local feedback rules interact with astrocytic domain organization and long-range glial signaling.

Beyond the regulation of externally driven network states, the model identified a distinct role for astrocytic signaling kinetics in shaping self-sustained activity following removal of external input. Importantly, our results distinguish two complementary roles of astrocytic signaling in network dynamics. Feedback topology primarily determines the qualitative state adopted by the neuronal network, including asynchronous, synchronous, and quiescent regimes, whereas astrocytic signaling kinetics regulate the temporal organization of self-sustained activity within these regimes. In particular, the kinetics of calcium and gliotransmitter signaling influenced both the frequency and persistence of Up states following removal of external stimulation. This separation between state control and temporal control suggests that astrocytes may regulate cortical dynamics at multiple computational levels: through pathway-specific modulation of network excitability and through slower control of the timing and persistence of collective activity.

Overall, these simulations support a mechanistic view in which the functional consequences of astrocytic signaling emerge from the interaction between cellular specificity, feedback topology, and signaling kinetics. Importantly, the present findings should be interpreted as model-based predictions rather than direct demonstrations of physiological mechanisms. The model predicts that selective manipulation of the neuronal populations driving or receiving astrocytic feedback should differentially affect cortical synchrony, excitation–inhibition balance, and the persistence of Up states. These predictions can be tested experimentally using cell-type-specific optogenetic or chemogenetic approaches, providing a direct route for evaluating the proposed neuron–astrocyte mechanisms in cortical circuits.

## References

Ali, O.B.K., Vidal, A., Grova, C., Benali, H., 2025. Dialogue mechanisms between astrocytic and neuronal networks: A whole-brain modelling approach. PLOS Computational Biology 21, e1012683.

Allen, N.J., Eroglu, C., 2017. Cell biology of astrocyte-synapse interactions. Neuron 96, 697–708.

Augusto-Oliveira, M., Arrifano, G.P., Takeda, P.Y., Lopes-Araújo, A., Santos-Sacramento, L., Anthony, D.C., Verkhratsky, A., Crespo-Lopez, M.E., 2020. Astroglia-specific contributions to the regulation of synapses, cognition and behaviour. Neuroscience & Biobehavioral Reviews 118, 331–357.

Barranca, V.J., Johnson, D.C., Moyher, J.L., Sauppe, J.P., Shkarayev, M.S., Kovačič, G., Cai, D., 2014. Dynamics of the exponential integrate-and-fire model with slow currents and adaptation. Journal of computational neuroscience 37, 161–180.

Blum Moyse, L., Berry, H., 2022. Modelling the modulation of cortical up-down state switching by astrocytes. PLoS Computational Biology 18, e1010296.

Breithausen, B., Kautzmann, S., Boehlen, A., Steinhäuser, C., Henneberger, C., 2020. Limited contribution of astroglial gap junction coupling to buffering of extracellular k+ in ca1 stratum radiatum. Glia 68, 918–931.

Brette, R., Gerstner, W., 2005. Adaptive exponential integrate-and-fire model as an effective description of neuronal activity. Journal of neurophysiology 94, 3637–3642.

Bushong, E.A., Martone, M.E., Jones, Y.Z., Ellisman, M.H., 2002. Protoplasmic astrocytes in ca1 stratum radiatum occupy separate anatomical domains. Journal of Neuroscience 22, 183–192.

Chen, Z., Yuan, Z., Yang, S., Zhu, Y., Xue, M., Zhang, J., Leng, L., 2023. Brain energy metabolism: astrocytes in neurodegenerative diseases. CNS neuroscience & therapeutics 29, 24–36.

Connors, B.W., Gutnick, M.J., 1990. Intrinsic firing patterns of diverse neocortical neurons. Trends in neurosciences 13, 99–104.

Cooper, M.L., Selles, M.C., Cammer, M., Redd, C., Gildea, H.K., Sall, J., Chiurri, K.E., Cheung, P., Wheeler, D.G., Saab, A.S., et al., 2026. Astrocytes connect specific brain regions through plastic networks. Nature, 1–9.

Destexhe, A., 2009. Self-sustained asynchronous irregular states and up– down states in thalamic, cortical and thalamocortical networks of non-linear integrate-and-fire neurons. Journal of computational neuroscience 27, 493–506.

Fisher, T.M., Liddelow, S.A., 2024. Emerging roles of astrocytes as immune effectors in the central nervous system. Trends in immunology 45, 824–836.

Garcia, D.W., Jacquir, S., 2024. Astrocyte-mediated neuronal irregularities and dynamics: the complexity of the tripartite synapse. Biological Cybernetics 118, 249–266. doi:10.1007/s00422-024-00994-z.

Garcia, D.W., Jacquir, S., 2025. From quiescence to self-sustained activity: How astrocytes reshape neural dynamics. Neuroscience 576, 182–198. URL: https://www.sciencedirect.com/science/article/pii/S0306452225002866, doi:10.1016/j.neuroscience.2025.04.009.

Goldman, J.S., Kusch, L., Yalcinkaya, B.H., Depannemaecker, D., Nghiem, T.A.E., Jirsa, V., Destexhe, A., 2020. Brain-scale emergence of slow-wave synchrony and highly responsive asynchronous states based on biologically realistic population models simulated in the virtual brain. BioRxiv, 2020–12.

Guthrie, P.B., Knappenberger, J., Segal, M., Bennett, M.V., Charles, A.C., Kater, S.B., 1999. Atp released from astrocytes mediates glial calcium waves. Journal of Neuroscience 19, 520–528.

Halassa, M.M., Fellin, T., Takano, H., Dong, J.H., Haydon, P.G., 2007. Synaptic islands defined by the territory of a single astrocyte. Journal of Neuroscience 27, 6473–6477.

Han, R.T., Kim, R.D., Molofsky, A.V., Liddelow, S.A., 2021. Astrocyte-immune cell interactions in physiology and pathology. Immunity 54, 211–224.

Kilgore, J.A., Kopsick, J.D., Ascoli, G.A., Adam, G.C., 2025. Biologically-informed excitatory and inhibitory ratio for robust spiking neural network training. Scientific reports 15, 24798.

Lee, H.G., Wheeler, M.A., Quintana, F.J., 2022. Function and therapeutic value of astrocytes in neurological diseases. Nature reviews Drug discovery 21, 339–358.

Lorenzo, J., Rico-Gallego, J.A., Binczak, S., Jacquir, S., 2025. Spiking neuron-astrocyte networks for image recognition. Neural Computation 37, 635–665.

Lorenzo, J., Vuillaume, R., Binczak, S., Jacquir, S., 2020. Spatiotemporal model of tripartite synapse with perinodal astrocytic process. Journal of computational neuroscience 48.

Manninen, T., Aćimović, J., Linne, M.L., 2023. Analysis of network models with neuron-astrocyte interactions. Neuroinformatics 21, 375–406.

Naud, R., Marcille, N., Clopath, C., Gerstner, W., 2008. Firing patterns in the adaptive exponential integrate-and-fire model. Biological cybernetics 99, 335–347.

Ogata, K., Kosaka, T., 2002. Structural and quantitative analysis of astrocytes in the mouse hippocampus. Neuroscience 113, 221–233.

Orthmann-Murphy, J.L., Abrams, C.K., Scherer, S.S., 2008. Gap junctions couple astrocytes and oligodendrocytes. Journal of Molecular Neuroscience 35, 101–116.

Oschmann, F., Berry, H., Obermayer, K., Lenk, K., 2018. From in silico astrocyte cell models to neuron-astrocyte network models: A review. Brain research bulletin 136, 76–84.

Postnov, D., Koreshkov, R., Brazhe, N.A., Brazhe, A.R., Sosnovtseva, O.V., 2009. Dynamical patterns of calcium signaling in a functional model of neuron–astrocyte networks. Journal of biological physics 35, 425–445.

Postnov, D., Ryazanova, L., Sosnovtseva, O., 2007. Functional modeling of neural–glial interaction. Biosystems 89, 84–91. doi: 10.1016/j.biosystems.2006.04.012. selected Papers presented at the 6th International Workshop on Neural Coding.

Rupareliya, V.P., Singh, A.A., Butt, A.M., Kumar, H., et al., 2023. The “molecular soldiers” of the cns: Astrocytes, a comprehensive review on their roles and molecular signatures. European Journal of Pharmacology 959, 176048.

Sanchez-Vives, M.V., Manasanch, A., Pigorini, A., Arena, A., Camassa, A., Juel, B.E., Porta, L.D., Capone, C., De Luca, C., De Bonis, G., et al., 2025. Multiscale dynamical characterization of cortical brain states: from synchrony to asynchrony. arXiv preprint arXiv:2510.05815 .

Santello, M., Volterra, A., 2009. Synaptic modulation by astrocytes via ca2+-dependent glutamate release. Neuroscience 158, 253–259.

Schummers, J., Yu, H., Sur, M., 2008. Tuned responses of astrocytes and their influence on hemodynamic signals in the visual cortex. Science 320, 1638–1643.

Soriano, J., Rodríguez Martínez, M., Tlusty, T., Moses, E., 2008. Development of input connections in neural cultures. Proceedings of the National Academy of Sciences 105, 13758–13763.

Stimberg, M., Brette, R., Goodman, D.F., 2019a. Brian 2, an intuitive and efficient neural simulator. elife 8, e47314.

Stimberg, M., Goodman, D.F., Brette, R., Pittà, M.D., 2019b. Modeling neuron–glia interactions with the brian 2 simulator, in: Computational glioscience. Springer, pp. 471–505.

Stogsdill, J.A., Harwell, C.C., Goldman, S.A., 2023. Astrocytes as master modulators of neural networks: Synaptic functions and disease-associated dysfunction of astrocytes. Annals of the New York Academy of Sciences 1525, 41–60.

Tesler, F., Linne, M.L., Destexhe, A., 2023. Modeling the relationship between neuronal activity and the bold signal: contributions from astrocyte calcium dynamics. Scientific Reports 13, 6451.

Torao-Angosto, M., Manasanch, A., Mattia, M., Sanchez-Vives, M.V., 2021. Up and down states during slow oscillations in slow-wave sleep and different levels of anesthesia. Frontiers in systems neuroscience 15, 609645.

Touboul, J., Brette, R., 2008. Dynamics and bifurcations of the adaptive exponential integrate-and-fire model. Biological cybernetics 99, 319–334.

Valles, S.L., Singh, S.K., Campos-Campos, J., Colmena, C., Campo-Palacio, I., Alvarez-Gamez, K., Caballero, O., Jorda, A., 2023. Functions of astrocytes under normal conditions and after a brain disease. International Journal of Molecular Sciences 24, 8434.

Verkhratsky, A., Matteoli, M., Parpura, V., Mothet, J.P., Zorec, R., 2016. Astrocytes as secretory cells of the central nervous system: idiosyncrasies of vesicular secretion. The EMBO journal 35, 239–257.

Verkhratsky, A., Nedergaard, M., 2018. Physiology of astroglia. Physiological reviews 98, 239–389.

Vuillaume, R., Lorenzo, J., Binczak, S., Jacquir, S., 2021. A computational study on synaptic plasticity regulation and information processing in neuron-astrocyte networks. Neural Computation 33, 1970–1992.

Zerlaut, Y., Chemla, S., Chavane, F., Destexhe, A., 2018. Modeling mesoscopic cortical dynamics using a mean-field model of conductance-based networks of adaptive exponential integrate-and-fire neurons. Journal of computational neuroscience 44, 45–61.

Zhou, B., Zuo, Y.X., Jiang, R.T., 2019. Astrocyte morphology: Diversity, plasticity, and role in neurological diseases. CNS neuroscience & therapeutics 25, 665–673.

